# Humans trade-off whole-body energy cost to avoid overburdening muscles while walking

**DOI:** 10.1101/2022.03.16.484670

**Authors:** Kirsty A. McDonald, Joseph P. Cusumano, Andrew Hieronymi, Jonas Rubenson

## Abstract

Metabolic cost minimization is thought to underscore the neural control of locomotion. Yet, avoiding high muscle activation, a cause of fatigue, often outperforms energy minimization in computational predictions of human gait. Discerning the relative importance of these criteria in human walking has proved elusive, in part, because they have not been empirically decoupled. Here we explicitly decouple whole-body metabolic cost and ‘fatigue-like’ muscle activation costs (estimated from electromyography) by pitting them against one another using two distinct gait tasks. When experiencing these competing costs, participants (*N*=10) chose the task that avoided overburdening muscles (fatigue avoidance) at the expense of higher metabolic power (p<0.05). Muscle volume-normalized activation more closely models energy use and was also minimized by the participants’ decision (p<0.05), demonstrating that muscle activation was, at best, an inaccurate signal for metabolic energy. Energy minimization was only observed when there was no adverse effect on muscle activation costs. By decoupling whole-body metabolic and muscle activation costs, we provide among the first empirical evidence of humans embracing non-energetic optimality in favor of a clearly defined neuromuscular objective. This finding indicates that local muscle fatigue and effort may well be key factors dictating human walking behavior and its evolution.

## Introduction

Humans often move in ways that save metabolic energy. For example, preferred walking speed (1–5), step frequency (6–12), step width (13) and arm swing (14,15) are classic examples known to minimize the metabolic cost of locomotion. More recently, it has been demonstrated that humans continue to exhibit energy-favoring behaviors, even after normal gait mechanics are disrupted (8,16,17). Given the consistency of these observations in human locomotion, and the selective advantage that they purportedly bestow, it is argued that whole-body energy minimization is a central factor dictating human gait behavior and its evolution (18,19).

However, whether metabolic cost minimization is itself the principal, overriding, optimality criterion during gait remains a matter of debate. For example, it has been proposed that other factors, such as fatigue avoidance, may be prioritized in the control scheme of locomotion (20–22), with low metabolic cost arising as a byproduct. Similarly, others suggest that economical human gait arises not from minimization of metabolic cost alone, but rather via the control of metabolic energy in conjunction with several additional optimality criteria (23,24).

Of the possible non-energetic optimality criteria, a compelling argument can be made for a muscle activation-based control signal in human locomotion. Several groups have developed successful simulations of human locomotion based on minimizing total muscle activation. This approach has proven particularly powerful in predicting realistic locomotor mechanics and energetics, often outperforming simulations based on minimizing total energy cost alone (21,22,25,26). For example, Miller et al. (21) discovered that more realistic gait biomechanics and whole-body metabolic cost could be predicted by minimizing muscle activation compared to metabolic cost itself, despite the longstanding view that humans consistently prioritize energy optimizing gaits (5,11). Likewise, Falisse et al. (23), and more recently, Veerkamp et al. (24), demonstrated that the inclusion of muscle activation in a multi-objective performance criterion is essential for generating the most physiologically realistic simulations of human gait. There also exists experimental data that point to muscle activation as a control signal in human locomotion. For example, the metabolic cost of walking on inclines of up to 10% can be predicted using muscle activation data (27). Preferred stride frequencies (28,29) and the walk-run transition (30–32)—a fundamental locomotor behavior that is not always tracked by whole-body energy cost (20)—can also be predicted by a muscle activation minimizing criterion. Similarly, self-selected cadence in cycling has been found to coincide with low muscle activity (33), more so than with minimization of metabolic energy expenditure (34,35). Together, these simulation and experimental data indicate that muscle activation may indeed be among the primary optimality criteria governing human gait.

On closer examination of the aforementioned studies, the activation cost functions that have been most successful in predicting gait mechanics and energetics (often better than energy optimization) penalize high activations in any given muscle. These cost functions minimize a weighted average of individual muscle activations (*A*) raised to an integer power (e.g., *A*^2^, *A*^3^ *etc*.), where the activation of a muscle is defined as the fraction (between 0 and 1) of the total muscle fibers activated (21,22). Other successful simulations of muscle control have used a similar, but more extreme formulation, whereby the maximum activation of any single muscle is minimized (the min-max model; (36)). As above, the resulting recruitment pattern is one that favors a uniform distribution of activations across muscles as opposed to high activations among a select few muscles—even in cases where the latter results in lower total energy consumption (22,26). This avoidance of high muscle activation that arises from non-uniform muscle recruitment has been classified as a fatigue-minimizing (endurance maximizing) strategy (22,37,38). Thus, the nervous system may prioritize a ‘division of labor’ across muscles that minimizes overburdening any given muscle, as opposed to a strict ‘economy of labor’. In this scenario, economical gait may arise not as the principal optimization criterion, but rather secondary to a control paradigm favoring even muscle recruitment (22).

Establishing the mechanism(s) driving locomotor behavior is key to understanding the control of human gait and the resulting energetics. Previous investigations have largely been limited to simulation studies that assign objective functions (optimality criteria) *a priori* and subsequently assess their efficacy based on their ability to reproduce a high number of human gait features. To the best of our knowledge, no study has empirically addressed whether whole-body energy use or fatigue-like muscle activation costs (e.g., sum of *A*^2^; (22)) are prioritized during locomotion. Therefore, we designed an experiment to pit fatigue-like muscle activation costs against the total metabolic cost during walking by disrupting the distribution of activation across the lower limb muscles. Importantly, in this design, moving with low energy expenditure while simultaneously favoring a low muscle activation cost was unfeasible, therefore providing an opportunity to empirically test whether one criterion was favored over the other.

## Materials and methods

### Participants

Ten healthy adults (5F, 5M; age 31.1±7.2 years; mass 69.6±11.1 kg; height 1.70±0.07 m; mean±SD) participated in this study after providing informed consent. The study was approved by the Institutional Review Board at The Pennsylvania State University and received reciprocal approval by the Human Research Ethics Committee at the University of Western Australia. Volunteers were required to be free from medical conditions that could make moderate-intensity exercise unsafe (e.g., heart, respiratory and/or musculoskeletal conditions).

### Experimental design overview

The primary objective of our experimental design was to create two competing gait conditions that the participants had to choose between. In our case, the two competing conditions were either i) moving with a low whole-body metabolic cost or ii) moving with a low activation cost (see below for cost functions). Importantly, in selecting between this Competing-Cost-Pair, one condition had to be traded off against the other; in other words, the metabolic cost and activation cost could not be simultaneously reduced. Through pilot testing, two locomotor conditions were established to meet this experimental design goal. The first condition, crouch walking on a level treadmill, was determined to elicit a moderate metabolic cost and a high activation cost due to the uneven distribution of muscle recruitment across lower limb muscles. The second condition, incline walking, included five levels of treadmill incline that ranged from 0-24% and that predominantly taxed metabolic cost. As the incline level was increased, a Competing-Cost-Pair was determined by identifying the incline level that elicited a metabolic cost exceeding that of crouch walking, while at the same time having a lower muscle activation cost (see Fig. 1).

**Figure 1.**
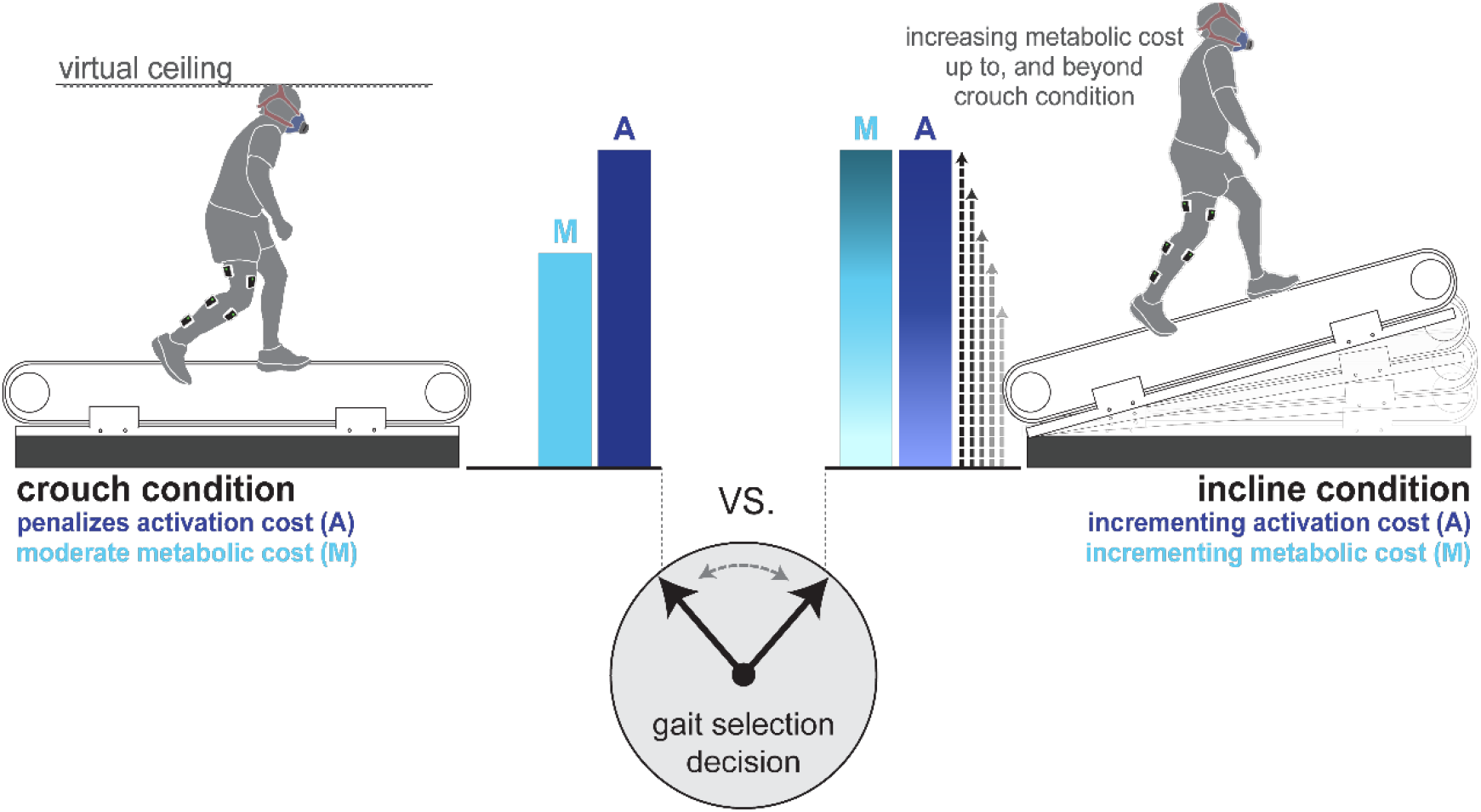
Experimental design. Participants were asked to select between crouch walking, which moderately affects metabolic cost and penalizes activation cost, and a series of incline levels that incrementally increase both metabolic (M) and activation (A) costs. A Competing-Cost-Pairs was established when participants walked on an incline level that incurred a higher metabolic cost but provided an activation cost advantage relative to the crouch condition. A virtual ceiling and the participant’s silhouette, displayed on a screen anterior to the treadmill, provided feedback regarding crouch walking task performance.

To establish the Competing-Cost-Pairs, five independent comparison trials were performed in a random order between flat (i.e., level treadmill) crouch walking and upright walking on one of the five incline levels (0%, 6%, 12%, 18% or 24%; Fig. 1). For each comparison, participants were asked to explore both crouch walking and upright walking on the assigned incline for three minutes. After this time, they selected their preferred state (crouch or upright incline walking), in which they then walked for an additional five minutes. The steepest incline level that was selected over crouch walking was referred to as the pre-transition incline. The first incline level for which the crouch walking was preferred was referred to as the post-transition incline (data were collected at the end of the five comparison trials for all incline levels that were rejected in favor of crouch walking). For example, if a participant chose the incline for 0-18% grades, and crouch walking at 24%, their pre-transition incline was defined as 18% and their post-transition incline was defined as 24%.

### Crouch walking

Crouch walking was achieved by having participants walk on a level treadmill (0% incline) while avoiding contact between their body and a virtual ceiling, with the specific instruction to maintain their trunk and head in a normal, upright walking posture (i.e., crouched, but not hunched or with lowered head). A motion sensing device (Xbox Kinect, Microsoft, WA, USA), situated behind the participant, was programmed to detect the participant’s body shape and produce a two-dimensional body silhouette. This silhouette was displayed on a large screen in front of the treadmill together with the virtual ceiling projection (see Supplementary Material). The ceiling height was programed to equal 93% of the participant’s standing height. This height was established in pilot testing to result in a crouch posture that elicited the desired increases in our metric of activation cost (see below). During periods of silhouette-ceiling collision, the outline of the silhouette was filled blue to alert the participant of contact. The number and duration of collisions was recorded for each trial. The real-time motion-tracking and visual feedback was implemented in a custom program developed with the openFrameworks open-source C++ toolkit.

At the beginning of each experiment, participants were provided a crouch walking familiarization trial. Participants walked at a set speed of 1.0 m s^−1^, crouching below the virtual ceiling. After three minutes of crouch walking with unlimited silhouette-ceiling collisions, the participant was informed that, to end the familiarization session, they must perform one minute of walking with no recorded collisions.

All participants achieved the one-minute benchmark and progressed to the next phase of data collection, in which their preferred crouch walking speed (PWS_C/W_; m s^−1^) was assessed in accordance with the method outlined by Dingwell and Marin (39). For all crouch and incline walking trials described below, the treadmill belt speed was set to the PWS_C/W_ so as not to disadvantage metabolic energy use in the crouch condition. Although it was not empirically verified that the PWS_C/W_ generated the lowest metabolic cost of all possible speeds for crouch walking (this was outside the scope of the current experiment), we speculated our protocol may provide a metabolic advantage in the crouch condition compared to other possible speeds (for example, the upright preferred walking speed). This was based on previous observations that humans will self-select walking speeds that produce the lowest metabolic cost for a given distance travelled (2,5,40).

### Comparison trials

Following PWS_C/W_ determination, a five-minute trial of crouch walking was recorded (initial crouch). In a randomized order, five levels of upright incline walking (on treadmill inclines of 0%, 6%, 12%, 18% and 24%) were then pitted against crouch walking, one by one. Each comparison trial (crouch versus a single incline level) commenced with 30 seconds of crouch walking and 30 seconds of incline walking (the order of which was reversed for consecutive comparison trials), followed by a further two minutes where the participant was free to explore both the crouch and incline. During the two-minute exploratory period, transition between the crouch and incline walking was verbally requested by the participant and manually initiated by the investigator, taking less than 10 seconds to implement.

At the conclusion of the exploratory period, the participant verbally expressed the condition (crouch or incline) in which they would prefer to walk for the following five minutes. Their preference was then imposed for a five-minute period, during which metabolic, surface electromyography (EMG), and ground reaction force (GRF) data were collected (details below). On discussions with participants at the conclusion of the experiment, it was confirmed that, together, the one minute (total) of imposed walking in the two conditions and the two minutes of free exploration was adequate for all participants to conclusively decide on a preferred condition.

Once crouch walking had been compared to the five incline levels, an additional five-minute crouch walking trial was recollected to account for fatigue/familiarization effects (final crouch). Any outstanding incline levels for which data was not collected as part of the comparison trials (because crouch walking was preferred) were then completed at PWS_C/W_, each for five minutes (randomized order), to provide insight into the participant’s motivation for transitioning (rejecting these inclines). To minimize any persistent muscle fatigue effects, participants commenced each consecutive trial when they verbally agreed that they felt they had returned to a ‘baseline (resting) level of exertion’. When describing the protocol, participants were made aware that they may be asked to complete at most 12 walking trials. However, the nature of the trials following the comparison trials were not revealed to the participants to ensure it did not influence their prior decisions.

### Whole-body energetics and electromyography measurements

Breath-by-breath oxygen uptake and carbon dioxide production were sampled using a portable metabolic system (Cosmed K4b^2^, Rome, Italy). A portable metabolic device was necessary to facilitate adherence to the required crouching posture and free regulation of body position on the treadmill. The final minute of data was used to compute a steady-state mass-specific metabolic power (*C_met,P_*; W kg^−1^) for each trial (details below). Surface EMG recordings were made synchronously from gluteus maximus (Gmax), biceps femoris (BF), rectus femoris (RF), vastus medialis (VM), medial gastrocnemius (MG), soleus (SOL) and tibialis anterior (TA) of the right leg, using silver-tipped surface electrodes placed according to SENIAM guidelines (41). Signals were recorded using a Bortec Octopus AMT-8 system (Calgary, Canada). Twenty strides of EMG data were collected at 2000 Hz. Ground reaction forces from the custom-built instrumented treadmill were also obtained (again sampled at 2000 Hz) to determine foot contacts and were later used to crop the EMG data into strides during data processing. Both EMG and ground reaction force data were obtained via a Cambridge Electronic Design data acquisition unit (Cambridge, UK). Metabolic, EMG and GRF data were collected during the initial and final crouch trials, all comparison trials where the incline was selected, and any outstanding incline trials (thus, there were seven total recording trials).

### Data processing

Data processing was performed in MATLAB 2018a (MathWorks, Natick, USA) using in-built functions and custom scripts.

#### Metabolic power (*C_met,P_*)

The rate of oxygen uptake 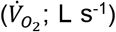 and carbon dioxide production 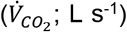 were averaged from the final minute of each trial and the following stoichiometric equation was applied to determine metabolic power (42,43):

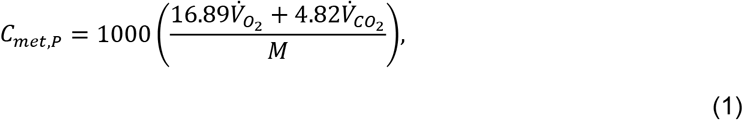

where *M* represents participant body mass. Because speed was constant across trials, we did not convert to a cost per distance traveled.

#### Muscle activation

Two participants’ activation data were eliminated from further analyses due to notable and persistent signal artefact in one of their processed electromyograms. For the remaining participants (*N*=8), the raw EMG data were DC offset, band-pass filtered (20-350 Hz) to remove any movement artefact and high frequency noise, and full-wave rectified. A linear envelope was applied to the rectified data using a low-pass filter with 6 Hz cut-off frequency (*EMG_env_*). All data filtering utilized fourth-order Butterworth underdamped filters (44). For each of the seven trials for which data was collected, five strides (*j* = 1,2, … 5) of *EMG_env_* data per muscle (*i* = 1,2, … 7) were individually integrated to produce scalar muscle activation values for each muscle, for each stride (*a_ij_*) as below:

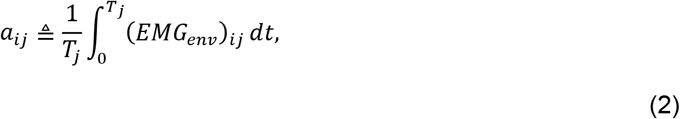

where (*EMG_env_*)*_ij_* represents the processed linear envelop of the *i^th^* muscle for the *j^th^* stride, 0 is the time at the initial contact of the right foot and *T_j_* is the stride time at the consecutive contact of the same (ipsilateral) foot determined from force plate thresholds (2% of peak signal). Thus, our expression for the muscle activation represents a combined effect of the number of fibers active within the muscles (the magnitude of the *EMG_env_* integral), as well as the rate at which they are recruited during continuous walking (the factor of 1/*T_j_*). This expression is similar to the fatigue activation cost function in Miller et al. (21).

A normalized (unitless) activation (*A_ij_*) was computed for each muscle, for each stride by normalizing by the average value obtained from the five strides of the 0% incline trial:

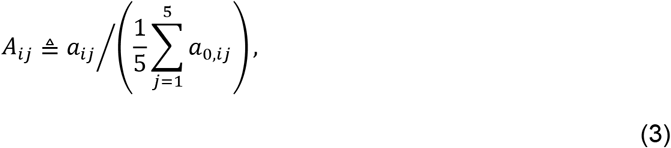

where *a_0_* is the activation integral of the 0% incline trial.

#### Activation cost functions

We patterned our activation cost functions on those presented by Ackermann and van den Bogert (22), which are based on a weighted average of muscle activations raised to a power *p*:

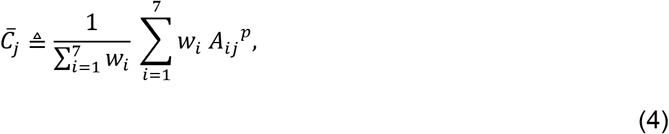

where 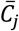 represents a generic cost for step *j* expressed as a weighted average over 7 muscles, *A_ij_* is as in Eq. (3), *p* is the exponent, and *w_i_* is the weighting factor for the *i^th^* muscle.

Using the general form of Eq. (4), equal muscle weights (*w_i_* = 1) and an exponent *p* > 1 leads to ‘fatigue-like’ activation cost functions that penalized large, fatigue-inducing, muscle activations irrespective of the size of the muscle (21,22). We used two such fatigue-like cost functions:

**a) Mean of squared activations 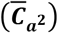; *w_i_* = 1 and *p* = 2.** For each trial, a total muscle activation cost was then computed as an average of the per-step cost of Eq. (4) across five strides:

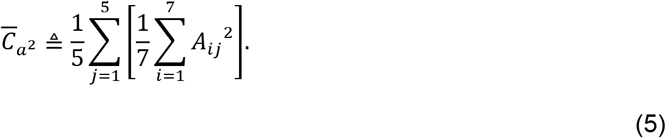

**b) Maximal muscle activation 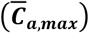; *w_i_* = 1 and *p* → ∞.** In the limit as *p* → ∞, only the maximally activated muscle contributes to the cost. This was then averaged across five strides to yield:

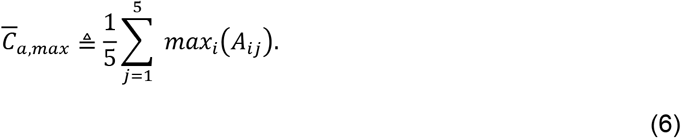

By adopting the cost functions, **(a)** and **(b)**, we capture a range of fatigue-like activation cost formulations, with the latter most strongly penalizing high activation in the individual muscles studied.

In contrast, when muscle volume weighting is used together with a low exponent, Eq. (4) is classified as an ‘effort-like’ cost (22). We included the strongest effort-like cost function, where muscle volume weights (*vol_i_*) are based on physiological muscle volumes and the exponent *p* = 1. This cost function does not penalize an uneven distribution of muscle activation as strongly as in Eqs. (5) and (6). Rather, it takes into consideration active muscle volume, a key determinant of muscle energy expenditure (45,46), by weighting the activations by muscle size. Therefore, unlike the fatigue-like cost functions, this cost is more sensitive to the activation of large muscles (22). This cost was included to assess the possibility that muscle activation serves as a proxy for metabolic energy expenditure (47).

**c) Volume-weighted muscle activations 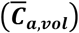; *w_i_* = *vol_i_* and *p* = 1**. The normalized activation rate (*A_ij_*) for each muscle was weighted according to the muscle volumes (*vol_i_*) documented in the supplementary material of Handsfield et al. (48). These weightings were 0.33, 0.08, 0.10, 0.17, 0.10, 0.17 and 0.05 for Gmax, BF, RF, VM, MG, SOL, and TA, respectively, and represent each muscle’s volume as a proportion of the sum of all seven muscles. The volume-weighted average of the seven normalized activation rates was computed using the general form of Eq. (4), and this value was again averaged across five strides:

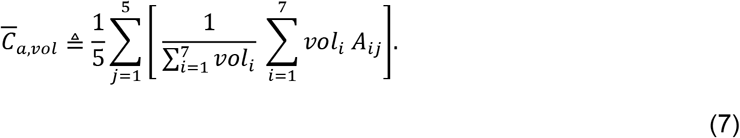

### Data analysis

For each metric described above, five values were initially analyzed; mean crouch (the mean value of the initial crouch and final crouch trials), initial crouch, final crouch, pre-transition incline (the steepest incline level which was selected over crouch walking) and post-transition incline (the lowest incline level that was rejected in favor of crouch walking). Whether the mean, initial or final crouch data were compared to the pre- and post-transition inclines had no bearing on the interpretation of the results discussed herein. Thus, the mean crouch data is represented by all future references to the crouch walking data.

#### Competing-Cost-Pairs

Each participant’s mean crouch walking data was compared to the pre-transition incline to determine whether a Competing-Cost-Pair had been established (i.e., the two trials were designed to provide mutually exclusive *C_met,P_*, and 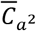 advantages). We accepted a Competing-Cost-Pair when *C_met,P_*, decreased betwe en the p re-t ransition incli ne and c rouc h walki ng trials, and 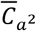 increased. The *C_met,P_*, and 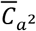 advantages were as sesse d because our aim was to compare metabolic and fatigue-like muscle activation costs, and 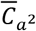 represents th e le ss ex treme (least biased) of our fatigue-like cost functions. A secondary analysis was then performed to determine whether Competing-Cost-Pairs were also present between *C_met,P_*, and 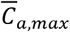, and *C_met,P_*, and 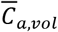.

### Statistical analyses

Summary statistics (group mean±SD) were computed for crouch walking performance variables (number and duration of silhouette-ceiling collisions), and for key cost metrics (*C_met,P_*, 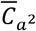, 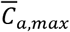 and 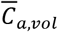) during the crouch and incline walking trials. Normality was assessed using the Shapiro-Wilk test. One-way repeated measures analysis of variance (ANOVA) tests were used to confirm main effects of the normally distributed incline data (0, 6, 12, 18, 24% inclines) for *C_met,P_*, 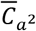, 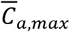 and 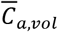.

For the crouch, pre-transition incline and post-transition incline comparisons, key cost metrics were each analyzed via a one-way repeated measures ANOVA (normally distributed) or Friedman test (non-normally distributed). *Post hoc* testing was then administered as paired-samples t-tests (normally distributed) or Wilcoxon tests (non-normally distributed). Results were compared against an adjusted alpha level (p=0.017; Bonferroni correction). All other analyses employed a standard 0.05 alpha level. In group-level analyses, all available data (*N*=10 for *C_met,P_*, and *N*=8 for 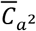, 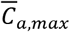, 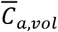) was used. Paired samples t-tests/Wilcoxon tests analyzed *C_met,P_*, 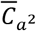, 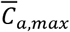 and 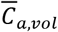 in the sub-sample of participants for whom a Competing-Cost-Pair was successfully established (*N*=7).

To discern whether the use of the initial, final or mean crouch data to represent crouch walking *C_met,P_*, 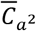, 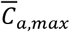, and 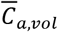 would make a difference to our interpretation of results, all relevant statistical tests (above) were performed three times: i) using the initial crouch data, ii) using the final crouch data, and iii) using the mean of the initial and final crouch data. No difference in interpretation was present and so mean crouch data is presented in the Results section below.

Any non-normally distributed data is denoted by a caret (^) in the Results section below. SPSS 26.0 (IBM, Chicago, USA) was used to perform all statistical analyses.

## Results

### Gait selections

In total, five decisions were made by our participants, i.e., one for each incline level. At low levels of incline (0-12%), all participants (*N*=10) chose the incline over the crouch. At 18% incline, the decision divided our sample; seven participants continued to select the incline, while three preferred the crouch (Fig. 2A). Those three participants also chose crouch over the 24% incline. The remaining seven participants all chose crouch walking over walking on the 24% incline.

**Figure 2.**
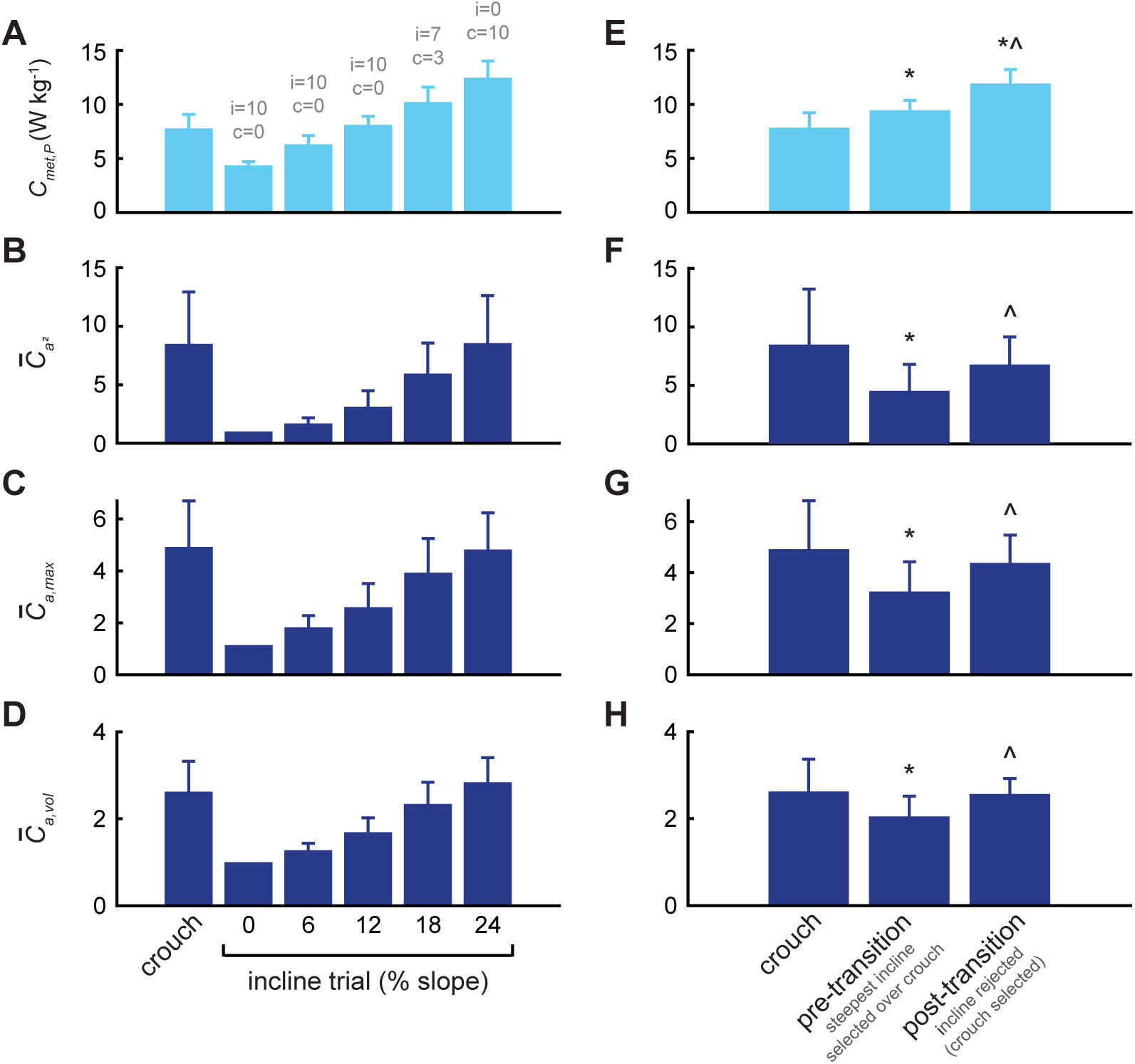
Walking data for the (A-D) mean (±SD) crouch trial and five levels of incline (from 0-24%) and (E-H) mean (±SD) crouch trial and pre- and post-transition inclines. (A, E) Metabolic power (*C_met,P_*; W kg^−1^; *N*=10), (B, F) mean of squared activations (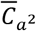; unitless; *N*=8), (C, G) maximal muscle activation (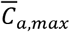; unitless; *N*=8), and (D, H) volume-weighted muscle activations (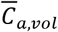; unitless; *N*=8). The number of participants that selected incline (i) or crouch (c) in each condition are indicated in (A) (grey text). *Significantly different (p<0.017) from crouch (E-H only); ^Significantly different (p<0.017) from pre-transition (E-H only).

The task performance of crouch walking was assessed by the total number and duration of silhouette-ceiling collisions during a single five-minute trial. To determine potential learning effects, these metrics were recorded at the beginning (initial crouch trial) and end (final crouch trial) of the data collection session. Total number (beginning: 1.2±1.5; end: 2.3±2.4) and duration (beginning: 0.11±0.15 seconds; end: 0.22±0.25 seconds) of collisions did not change significantly (p=0.131^ and p=0.075^, respectively). The mean PWS_C/W_ was 1.15 (±0.06) m s^−1^.

### Incline and crouch effects on activation and energy costs

All key cost metrics (*C_met,P_*, 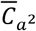, 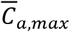 and 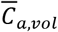) increased with increasing incline level (Fig. 2A-D). This was confirmed by one-way repeated measures ANOVAs, which demonstrated a main effect of incline for *C_met,P_* (p<0.001), 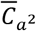 (p=0.001), 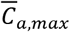 (p<0.001), and 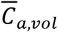 (p<0.001).

Crouch walking disrupted the distribution of muscle activation compared to upright walking, such that select muscles had proportionately higher activation values. In particular, Figure 3 highlights the increased activation requirements of knee extensor musculature (vastus medialis and rectus femoris) to facilitate the crouch gait, while the steepest incline gaits are achieved with a more even distribution of activation.

**Figure 3.**
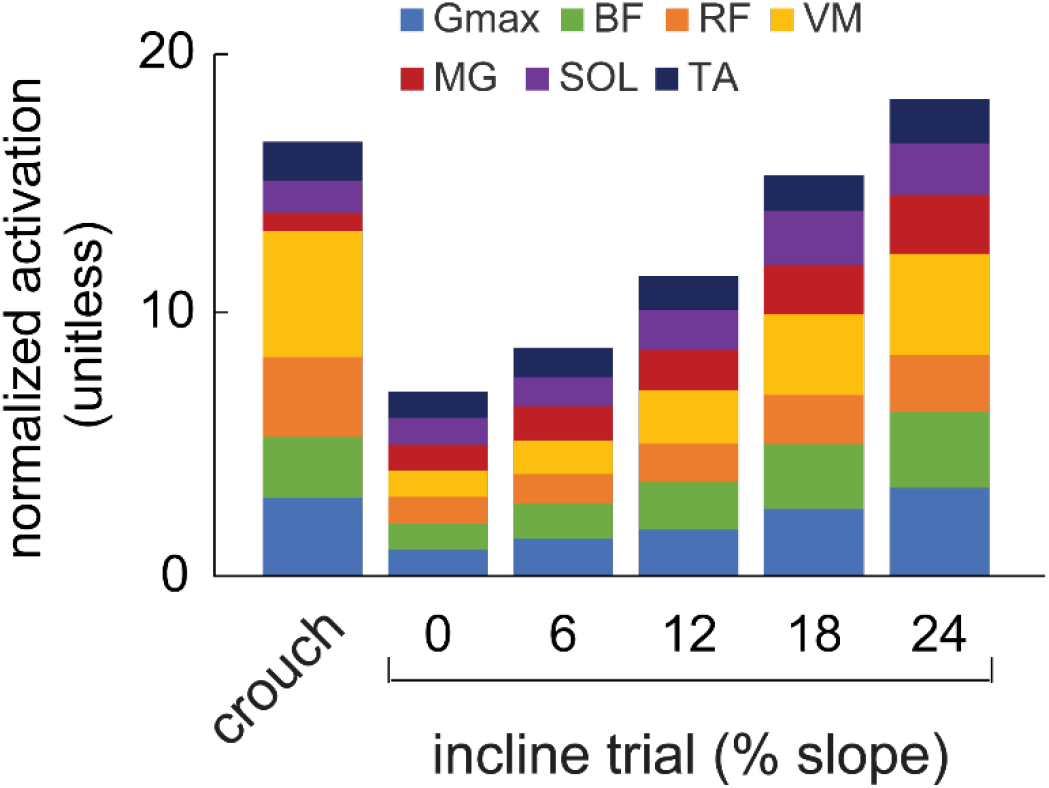
Average (*N*=8) average normalized activation costs during crouch walking (level treadmill) and incline walking on treadmill slopes of 0-24%. Seven lower limb muscles were analyzed; gluteus maximus (Gmax), biceps femoris (BF), rectus femoris (RF), vastus medialis (VM), medial gastrocnemius (MG), soleus (SOL) and tibialis anterior (TA). The unitless quantity *A_ij_* was established by integrating the linear envelope and expressing this as a rate (*a_ij_*), before normalizing data for each muscle to the average value obtained from the 0% incline trial. These normalized data were then averaged over five strides.

When statistical comparisons were made between the crouch, pre-transition incline and post-transition incline, the one-way repeated measures ANOVAs/Friedman tests demonstrated a main effect for *C_met,P_* (p<0.001, *N*=10), 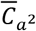 (p=0.008^; *N*=8), 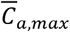 (p=0.001, *N*=8) and 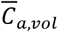 (p=0.003; *N*=8; Fig. 2E-H).

### Competing-Cost-Pairs

A Competing-Cost-Pair (i.e., two trials providing mutually exclusive *C_met,P_* and 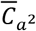 advantages) was effectively established for seven of the eight participants for whom we acquired a complete set of EMG data (Table 1; Fig. 4). Competing-Cost-Pairs could also be established for these participants between *C_met,P_*, and 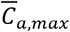, and *C_met,P_* and 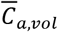 (Table 1).

**Table 1.**
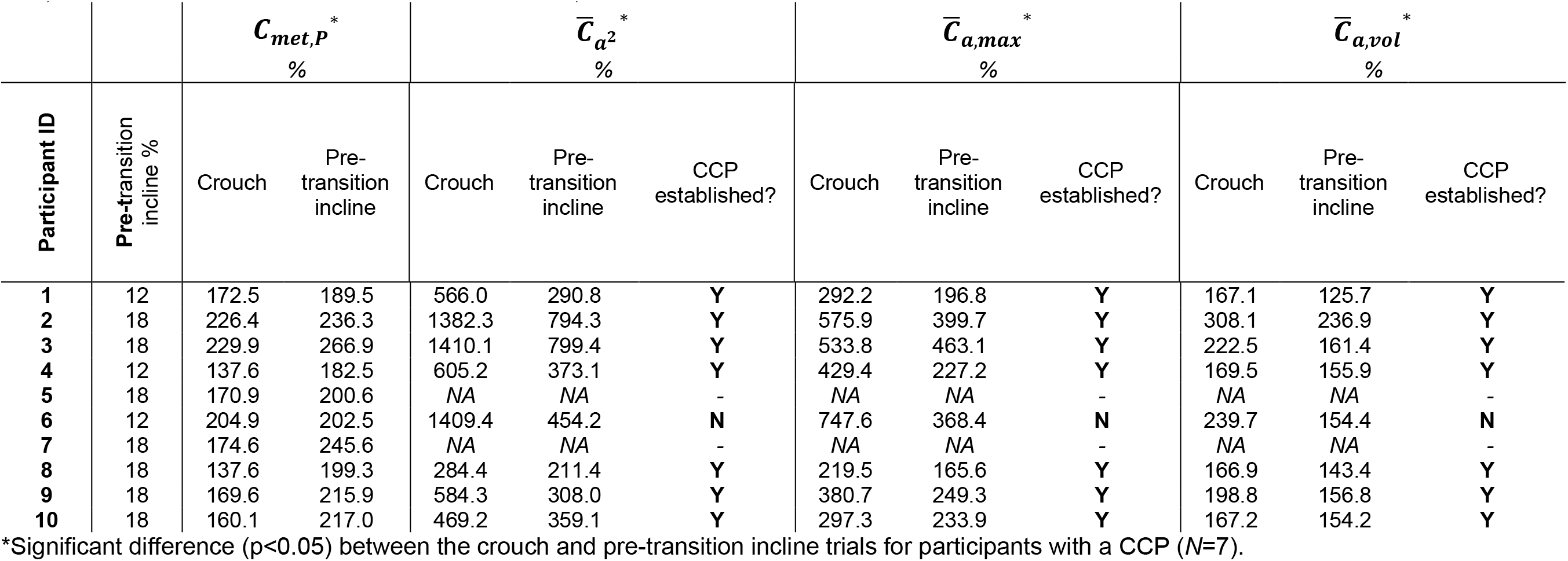
Competing-Cost-Pairs (CCPs) were established for seven of the eight participants for which this was feasible. The presence of a Competing-ost-Pair was determined by comparing the mean crouch and pre-transition incline’s percentage change from upright (normal) walking on a 0% cline for: i) metabolic power (*C_met,P_*) and the mean of the squared activations 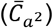, ii) *C_met,P_* and the maximal muscle activation 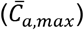, and iii) *C_met,P_* and the volume-weighted muscle activations 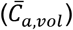. The pre-transition incline is defined as the steepest incline that was selected over crouch.

**Figure 4.**
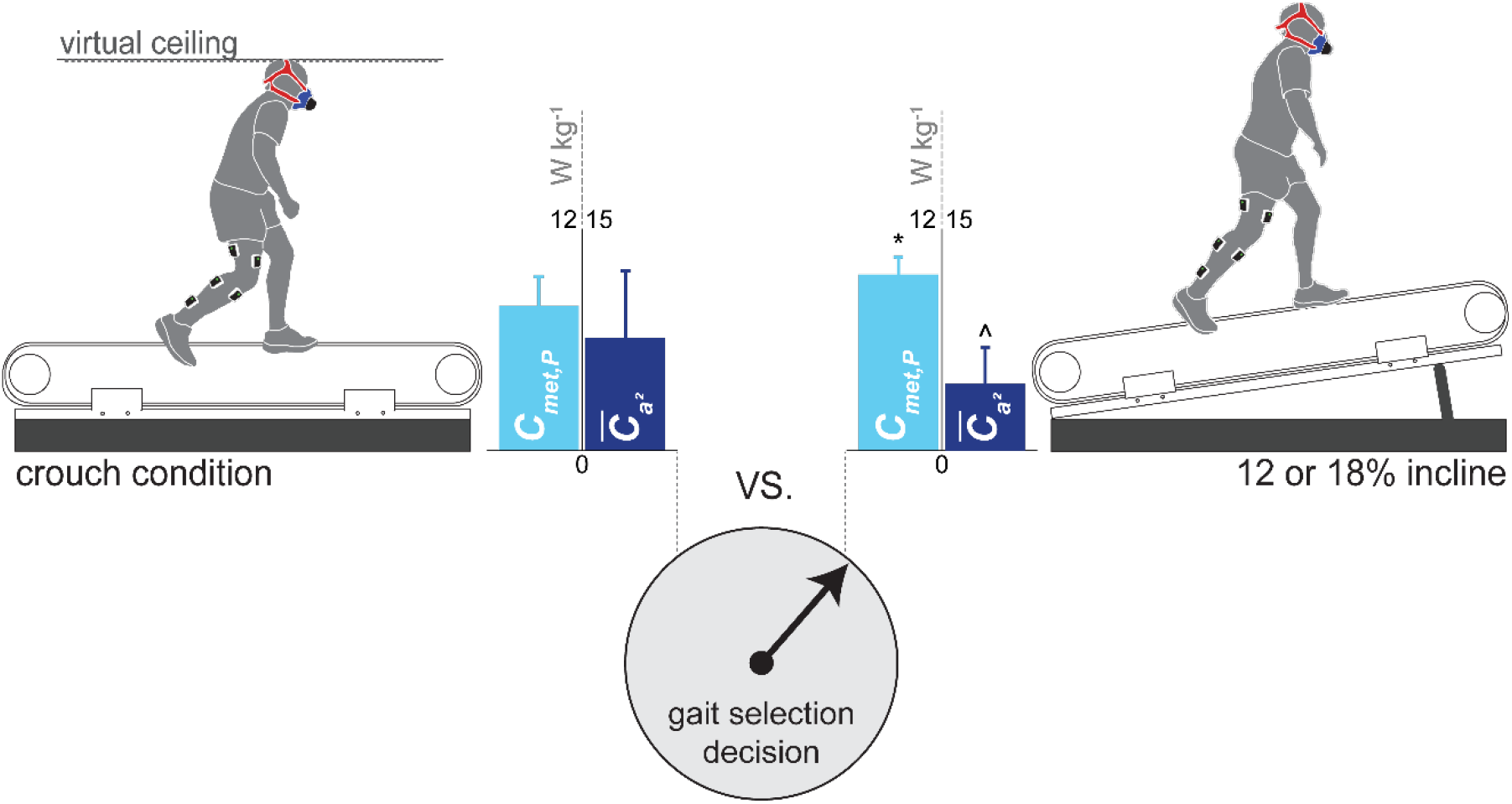
Competing-Cost-Pairs were established for seven of the eight participants for which this was feasible. All seven participants selected the incline condition that favored the mean of squared activations 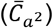 and penalized metabolic power (*C_met,P_*). In doing this, they rejected the crouch walking condition that favored metabolic power and penalized the mean of squared activations. Data is presented as the mean±SD. *Significantly different (p<0.05) from crouch *C_met,P_*; ^Significantly different (p<0.05) from crouch 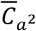.

All seven participants (1-4 and 8-10) chose incline walking over crouch walking and thus selected low muscle activation at the expense of a high *C_met,P_*. The average 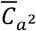 advantage in the Competing-Cost-Pairs was 66% (*N*=7; p=0.018^^^; range: 31 to 95%) and for 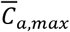 was 44% (*N*=7; p=0.003; range: 15 to 89%; Table 1). The average *C_met,P_* penalty was 19% (*N*=7; p=0.002; range: 4% to 31%; Table 1). Participant #6 had a *C_met,P_* that was nearly identical in the crouch and incline condition (1% difference) and therefore did not present a clear energetic advantage/disadvantage for gait selection (Table 1). However, Participant #6 achieved the group’s largest activation advantage by selecting the incline condition (210% reduction in 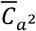; 103% reduction in 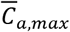). We also found an advantage in the effort-like activation cost 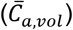 when the participants selected incline walking over crouch walking (Table 1). The average 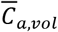 advantage in the Competing-Cost-Pairs was 23% (*N*=7; p=0.007; range: 8 to 38%).

In addition to the Competing-Cost-Pairs, prioritization objectives can be further assessed from comparing the group mean *C_met,P_* (*N*=10) and activation costs (*N*=8) of the crouch walking and pre- and post-transition incline conditions (Fig. 2E-H). Crouch walking incurred a *C_met,P_* of 7.84±1.38 W kg^−1^, which was significantly lower than both the pre-(9.46±0.91 W kg^−1^; p=0.007) and post-(11.92±1.31 W kg^−1^; p=0.005) transition incline levels, with the pre- and post-transition inclines being significantly different from one another (p=0.005; Fig. 2E). In contrast, the 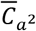 and 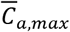 for the crouch walking were significantly greater (84%; p=0.012^^^ and 52% p=0.006, respectively) than the pre-transition incline (Fig. 2F-G). These data show, at a group level, a selection (pre-transition incline) for lower fatigue-like activation costs at the expense of higher *C_met,P_*. We also found that the 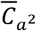 and 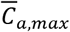 in crouch walking were statistically equivalent to the post-transition incline (p=0.263^^^ and p=0.214, respectively), indicating that the disadvantage in the fatigue-like activation costs was no longer present when crouch walking was finally selected (crouch walking versus post-transition incline walking). The effort-like activation metric (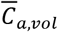 followed a similar pattern (Fig. 2H). 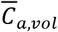 for crouch walking was significantly greater (27%) than for the pre-transition incline (p=0.003) but not for the post-transition incline (p=0.779).

## Discussion

Metabolic energy expenditure is widely regarded as a principal determinant of animal locomotor behavior (5,8,11,49). However, simulation studies of human locomotion suggest muscle activation may also be a key control parameter (21,22). It is difficult to discern how these criteria are weighted in the control scheme of locomotion due to their highly coupled nature. The current study utilized an experimental design that systematically pitted whole-body metabolic cost against muscle activation costs. In these competing conditions, global energy expenditure was clearly not prioritized, and our empirical data provide promising support for muscle activity prioritization.

### Energy expenditure is not always the principal objective in human locomotion

Humans have been observed to expend more energy than minimally required for both upper and lower limb movement tasks (40,50–56). These studies suggest that energy minimization may be task dependent. Perhaps the most well documented example of non-energetically optimal movement is self-selected cycling cadence (25,33–35). In the context of gait, Yandell and Zelik (56) show inconsistent prioritization of metabolic cost when participants used their preferred step frequency during barefoot walking. The authors propose this may be related to the discomfort/pain associated with an unshod gait (56). Some evidence also suggests humans do not prioritize metabolic cost minimization when moving in less stable environments, e.g., walking downhill (50) (but this remains inconclusive (57)) or walking with experimentally imposed asymmetrical leg lengths (40). Humans have also been observed to perform walking tasks in more effortful ways to eliminate a cognitive burden (51). Furthermore, there exist several studies of human movement in which immediate, independent detection and/or acute prioritization of the energetic minima is not evident (8,40,58,59). The costs driving gait selection in the aforementioned studies remain speculative. The current study is, in contrast, among the first to explicitly and experimentally demonstrate an alternative criterion, namely muscle activation, that is prioritized over metabolic cost in the control scheme of locomotion (at least under the conditions studied here).

While it is clear that a *C_met,P_* penalty occurs with the selection of incline walking in the Competing-Cost-Pairs (and in the pre-transition incline selection more generally; Fig. 2E), we do not suggest that energy minimization is unimportant to the control of walking, even in less common situations such as the crouch walking investigated in this study. Indeed, in the final incline condition (post transition incline), when the *C_met,P_* was the highest and the activation penalty of crouch walking was reduced or eliminated, participants opted to save energy by selecting crouch walking. Rather, our study supports the idea that locomotion tasks are regulated in a multi-objective manner that includes metabolic cost as one important, but not the sole, factor. Taken together, our data across all conditions suggest that a priority for energy minimization is not an inevitability, but also that as *C_met,P_* increases so might the relative weighting of *C_met,P_* in the control scheme of walking.

### Does a ‘division of muscle labor’ dictate gait control?

The non-volume-weighted exponential activation cost function, 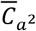, penalizes high (fatiguing) activations in any single muscle and thus favors an even muscle recruitment strategy (22). The hypothesis that gait is under a control scheme that minimizes muscle fatigue (or, more generally, overburdening of muscles) is not new, but has been mostly limited to simulation studies (21,22). To the best of our knowledge, the current study is the first to empirically observe a fatigue-like control scheme take precedence over a metabolic cost control scheme. Our finding that subjects protected their 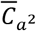 at the direct expense of *C_met,P_* was observed both in the individual participants’ Competing-Cost-Pair trials (Table 1), and also at a group (average) level (Fig. 2E-F).

The clear preference for protecting 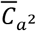 over energy cost suggests a strong control objective for maintaining an even muscle recruitment and thus avoiding high activations that can cause local muscle exhaustion (22). This is also evident from the more extreme cost function, 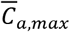, which showed that all participants’ gait selection in the Competing-Cost-Pairs protected their would-be maximally activated muscle, in line with a *min/max* cost function (36). This further suggests that the decision to avoid crouch walking may, in fact, have been specifically predicated on not exhausting the primary contributing muscle. Even if highly activated individual muscles represent a small fraction of the total muscle mass, they can have an important limit on performance because maximally activated muscles set the bottle neck for the endurance of the whole muscle system (22,60).

Further support for activation-driven gait selection comes from our observations of the post-transition incline gait (i.e., when crouch walking was finally chosen). When crouch walking is compared to the post-transition incline, we observe a diminished activation advantage. This can be seen in the group average data, where similar and not statistically different 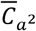,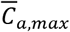 and 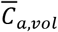 values were observed between conditions (Fig. 2F-H). Perhaps not only avoiding, but also selecting crouch walking, involved information from muscle activation. It may be that only when there is minimal/no activation disadvantage, do participants accept a *C_met,P_* advantage (whether this was selecting incline over crouch walking, or vice versa, selecting crouch walking over very steep incline walking). It is also worth pointing out that the high 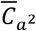 cost associated with crouch walking and the steepest incline walking resulted from different muscle recruitment patterns (Fig. 3); high activations in crouch walking are primarily in the vastus medialis (for 6/8 participants), while in the post-transition incline the muscles that exhibit the high activations are more variable between subjects.

Why should muscle activation be prioritized over energy cost? One answer may be that minimizing high activations that overburden or fatigue a given muscle will likely permit longer movement duration, a performance criterion that is regarded to be fundamental to the evolution of human bipedalism and with known ecological relevance (61–64). For instance, to effectively travel long distances in persistence hunting (regarded as a key feature of human evolution; 62) there is a clear selective advantage to resist fatigue. Muscle activation is a compelling control signal for sensing a division of labor among muscles and for avoiding overburdening individual muscles. It is thought that muscle effort can be readily sensed using central (e.g., via an efference copy) and peripheral (e.g., Group III and IV muscle afferents) mechanisms (65,66).

### Does muscle activation represent a proxy for metabolic energy use?

Notwithstanding the clear acceptance of a *C_met,P_* penalty, could it be possible that the nervous system was attempting to prioritize total metabolic energy expenditure using a proxy sensor, but failed? In their recent work, Wong and colleagues (67) concluded that blood-gas receptors are an unlikely sensor for whole-body locomotor energy expenditure and raised the possibility of a muscle-level sensor for metabolic cost. Muscle activation is a promising candidate for sensing energy cost because it is related to the fraction of the muscle that is metabolically active and because activation itself exacts a substantial metabolic cost in the form of calcium pumping (68–70). If activation serves as a proxy sensor for energy expenditure, the high volume-weighted activation signal associated with crouch walking may have ‘tricked’ the nervous system into registering a higher metabolic cost than that which was actually present. This could occur, in part, due to the inflated activation contribution of the large (high volume) knee-extensor muscles (Fig. 3). It is worth noting that in tasks where energy minimization is ubiquitously demonstrated (e.g., steady level walking), muscle activation and whole-body metabolic costs respond similarly to changing task parameters [e.g., step frequency; (11,28)], thus indicating that the muscle activations can, indeed, serve as a proxy for the energy consumption. This was, however, not the case in the present study. Both 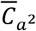, 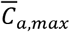, and 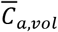 were disassociated from *C_met,P_*, with 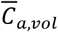 considered a proxy for energy use because it takes into account active muscle volume (22)—a key determinant of locomotor energy use (45,46,70,71).

Thus, if activation was used as a proxy for energy, it resulted in a misrepresentation of the relative energy cost of the crouch versus incline walking conditions. This scenario affects our interpretation of the energy optimization hypothesis in important ways. First, it would indicate that locomotor energy sensing is imperfect and may be cruder than previously thought. This implies that high-precision, rapid, energy sensing and subsequent gait control may be confined to common and predictable steady-state movements (e.g., steady level ground walking), and, in-turn, that energy optimization may not be particularly robust outside of a narrow range of conditions. Other experiments that have altered the mechanics of walking in novel ways have found that subjects often need guidance to discover energy minimizing movements (8,16). This suggests that while energy minimization may be achieved eventually, after sufficient motor learning has taken place, it may not be prioritized acutely. Whether trial-and-error learning can bring about versatile energy sensing across a myriad of natural conditions for human locomotion remains unclear.

Among the most interesting questions that remains is *how* the central nervous system uses muscle-level signaling to determine the physiological state of the body. For example, does the central nervous system achieve this by summing the activity of all muscles involved in a given task, weighted according to their size and tracked over a given distance or time period, and finally reference this against a threshold representing global (whole-body) effort or energy use (i.e., 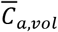)? Or, at the other end of the spectrum, is the control signal based on assessing individual muscle activation levels against thresholds unique to each actuator (i.e., 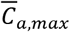)?

We posit a strategy assessing individual muscle activation levels against unique thresholds is a simpler control strategy; 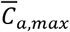 (and also, to a lesser extent, 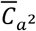) has a computational advantage, limiting the synthesis of information required by the central nervous system. Individual muscle activation thus presents a promising sensory modality when considering the credit-assignment problem (67,72), which refers to the direct or indirect nature of the signal(s) used by the central nervous system to steer gait selection behavior. Conversely, weighting and summing the individual activations requires the additional collation of information and is therefore a less direct signaling method for ascribing locomotor ‘cost’. The latter would therefore likely require greater computational effort and time.

We also do not, at present, have an effective way to rank the likelihood that one activation parameter is a more likely target of control than the other, although we do note that the pre-transition incline selection saw participants avoid greater 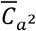 and 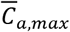 penalties than that associated with 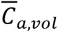 (84% and 52% versus 27%, respectively; *N*=8). Simulation studies that have established both mass- or volume-weighted activation and distribution cost functions also support this perspective, with more realistic gait patterns predicted by a non-volume-weighted or low volume-weighted muscle activation criterion (21,22). Irrespective of what optimality criterion activation minimization serves (e.g., a proxy for metabolic cost minimization, improved distribution of muscle effort, fatigue minimization, etc.), activation itself seems like a reasonable control target that can utilize relatively simple computational cost strategies.

### Alternate explanations for the observed gait selection

Despite the systematic prioritization of a gait with low muscle activation in the Competing-Cost-Pairs, we cannot rule out alternate hypotheses for the observed gait selection. For example, although speculative, cognitive factors, discomfort, pain, or other psychological motivators could be at the source of the selection of incline walking over crouch walking in the Competing-Cost-Pairs. Clearly delineating the role of discomfort versus activation *per se* in our observed gait selections is challenging because the two are related. In crouch walking, muscles with disproportionately high activations are prone to fatigue, which is known to cause discomfort (73). High muscle activations may also result in high local mechanical stresses that could trigger a pain response. This should not, however, detract from the observation that minimizing muscle activation, as opposed to metabolic energy, was a very strong predictor of movement selection in the present study. It is conceivable, for example, that high muscle activations are the proxy signals used in a neural computation of a ‘comfort’ cost. In this regard, the minimization of muscle activation in the Competing-Cost-Pairs could underlie a higher-level comfort objective.

### Limitations

The current study captured the activity of seven lower limb muscles that play a key role in human locomotion. Expanding upon previous empirical studies (47), we also include a weighting factor when determining the total activation cost. However, incorporating a greater selection of lower limb muscles and capturing trunk musculature contributions when computing activation costs would likely provide further insight into prioritization behaviors and may shed light on any differences in muscle recruitment strategies between participants. Furthermore, it is possible that the contributions of muscles that had low activation during our normalization trial (0% incline) were over-represented when costs were computed for other incline trials and/or the crouch walking trial. Taken together, it is possible that these limitations contributed to the variation seen in individual participant responses (Table 1).

While *C_met,P_* is susceptible to changes in both walking speed and step frequency (5,11), the latter was not imposed for any condition, and the former may have provided a metabolic advantage to crouch walking because all trials occurred at the preferred speed established for crouch walking. Yet, irrespective of this, participants still accepted a metabolic penalty through their pre-transition incline selection. It must be noted that whether one variable was closer to its physiological maximum than the other (e.g., 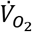 max versus maximal activation) was not measured in the current study. How close a variable is to its maximum may impact the priority for minimizing that variable. In this sense it is possible that selectively impairing other (more) important optimality criteria through energetic minimization will only occur in situations where a very high metabolic cost needs to be avoided. We acknowledge that the relative importance of activation and *C_met,P_* may also be different in steady-state level gait, and that it is likely task and condition dependent.

It is of course possible that other physiological signals exist whose distribution across muscles may parallel that of activation, for example afferent signals from muscle metabolism receptors (class III and IV afferents), or proprioceptive signals from muscle spindles and/or Golgi tendon organs. These may also be involved in the control scheme of walking and deserve further exploration together with muscle activation.

## Conclusion

Here, we provide among the first empirical evidence of locomoting humans accepting a metabolic cost penalty in favor of a predefined, alternative neuromuscular criterion. When choosing between crouch walking and walking on a series of increasing inclines, our healthy participants protected both fatigue-like and effort-like activation cost metrics. This observation is concomitant with activation cost prioritization in the control of steady-state walking (22) and running (21). Further research is required to determine whether muscle activation is used to estimate whole-body energy expenditure, or if local muscle costs are of greater importance in the control scheme of gait, independent of their relationship to energy costs *per se*.

## Acknowledgements and Funding Statement

The authors would like to thank Dr. Suzanne Cox and Mr. Brekke Green for data collection assistance and Drs. Brian Umberger, Ton van den Bogert, Lex Gidley, Jessica Selinger, Max Donelan, Greg Sawicki, Monica Daley, Young-Hui Chang, and Joseph Hamill for thoughtful discussions and/or feedback on earlier drafts of this manuscript. The authors also thank Mr. Randall McCullough for his technical support. This research received no specific funding from any funding agency in the public, commercial, or not-for-profit sectors.

---

**Figure S1.**
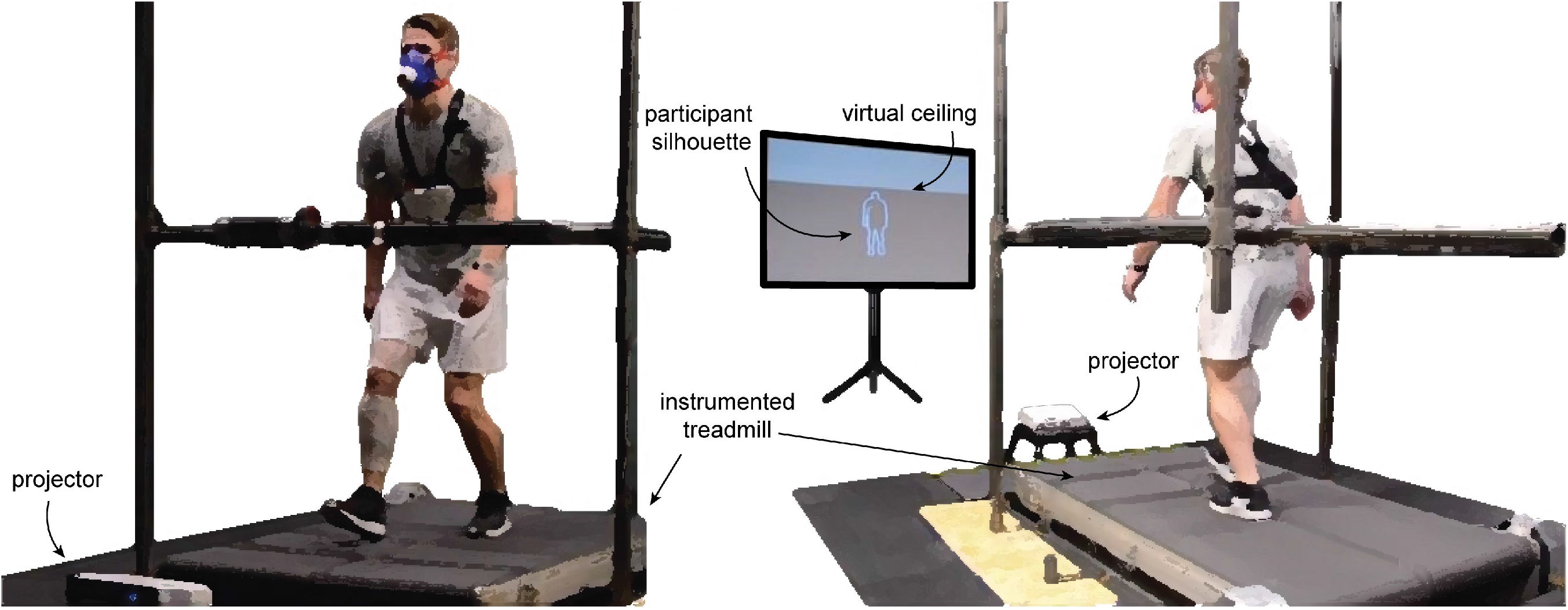
Experimental setup for crouch walking. Participants walked on a split-belt treadmill with their silhouette projected onto an anteriorly aced screen. Their objective was to avoid collisions between their silhouette and a virtual ceiling, which was set to 93% of their standing height. eal-time motion-tracking and visual feedback was implemented in a custom program developed with the openFrameworks open-source C++ toolkit, ing data from a motion sensing device (Xbox Kinect, Microsoft, WA, USA), situated behind the participant.

## References

1. Browning RC, Kram R. Energetic cost and preferred speed of walking in obese vs. normal weight women. Obesity. 2005;13(5):891–9.

2. Browning RC, Baker EA, Herron JA, Kram R. Effects of Obesity and Sex on the Energetic Cost and Preferred Speed of Walking. J Appl Physiol. 2006;100(2):390–8.

3. Holt KG, Jeng SF, Ratcliffe R, Hamill J. Energetic cost and stability during human walking at the preferred stride frequency. J Mot Behav. 1995;27(2):164–78.

4. Willis WT, Ganley KJ, Herman RM. Fuel oxidation during human walking. Metabolism. 2005;54(6):793–9.

5. Ralston HJ. Energy-speed relation and optimal speed during level walking. Eur J Appl Physiol Occup Physiol. 1958;17(4):277–83.

6. Minetti AE, Capelli C, Zamparo P, Di Prampero PE, Saibene F. Effects of stride frequency on mechanical power and energy expenditure of walking. Med Sci Sport Exerc. 1995;27(8):1194–202.

7. Molen NH, Rozendal RH, Boon W. Graphic representation of the relationship between oxygen-consumption and characteristics of normal gait of the human male. Proc K Ned Akad van Wet. 1971;75(4):305–14.

8. Selinger JC, O’Connor SM, Wong JD, Donelan JM. Humans can continuously optimize energetic cost during walking. Curr Biol. 2015;25(18):2452–6.

9. Snaterse M, Ton R, Kuo AD, Donelan JM. Distinct fast and slow processes contribute to the selection of preferred step frequency during human walking. J Appl Physiol. 2011;110(6):1682–90.

10. Umberger BR, Martin PE. Mechanical power and efficiency of level walking with different stride rates. J Exp Biol. 2007;210(18):3255–65.

11. Zarrugh MY, Todd FN, Ralston HJ. Optimization of energy expenditure during level walking. Eur J Appl Physiol Occup Physiol. 1974;33(4):293–306.

12. Zarrugh MY, Radcliffe CW. Predicting Metabolic Cost of Level Walking. Eur J Appl Physiol Occup Physiol. 1978;38(3):215–23.

13. Donelan JM, Kram R, Kuo AD. Mechanical and metabolic determinants of the preferred step width in human walking. Proc R Soc B. 2001;268(1480):1985–92.

14. De Graaf ML, Hubert J, Houdijk H, Bruijn SM. Influence of arm swing on cost of transport during walking. Biol Open. 2019;8(6).

15. Ortega JD, Fehlman LA, Farley CT. Effects of aging and arm swing on the metabolic cost of stability in human walking. Journal of biomechanics, 41(16), 3303–3308. J Biomech. 2008;41(16):3303-8.

16. Abram SJ, Selinger JC, Donelan JM. Energy optimization is a major objective in the real-time control of step width in human walking. J Biomech. 2019;91:85–91.

17. Roemmich RT, Leech KA, Gonzalez AJ, Bastian AJ. Trading symmetry for energy cost during walking in healthy adults and persons poststroke. Neurorehabil Neural Repair. 2019;33(8):602–13.

18. Wang WJ, Crompton RH, Li Y, Gunther MM. Energy transformation during erect and ‘bent-hip, bent-knee’walking by humans with implications for the evolution of bipedalism. J Hum Evol. 2003;44(5):563–79.

19. Rodman PS, McHenry HM. Bioenergetics and the origin of hominid bipedalism. Am J Phys Anthropol. 1980;52(1):103–6.

20. Hreljac A. Preferred and energetically optimal gait transition speeds in human locomotion. Med Sci Sport Exerc. 1993;25(10):1158–62.

21. Miller RH, Umberger BR, Hamill J, Caldwell GE. Evaluation of the minimum energy hypothesis and other potential optimality criteria for human running. Proc R Soc B. 2012;279(1733):1498–505.

22. Ackermann M, van den Bogert AJ. Optimality principles for model-based prediction of human gait. J Biomech. 2010;43(6):1055–60.

23. Falisse A, Serrancolí G, Dembia CL, Gillis J, Jonkers I, De Groote F. Rapid predictive simulations with complex musculoskeletal models suggest that diverse healthy and pathological human gaits can emerge from similar control strategies. J R Soc Interface. 2019;16(157):20190402.

24. Veerkamp K, Waterval NFJ, Geijtenbeek T, Carty CP, Lloyd DG, Harlaar J, et al. Evaluating cost function criteria in predicting healthy gait. J Biomech. 2021;110530.

25. Gidley AD, Marsh AP, Umberger BR. Performance criteria for generating predictive optimal control simulations of bicycle pedaling. Comput Methods Biomech Biomed Engin. 2019;22(1):11–20.

26. Song S, Geyer H. Predictive neuromechanical simulations indicate why walking performance declines with ageing. J Physiol. 2018;596(7):1199–210.

27. Silder A, Besier T, Delp SL. Predicting the metabolic cost of incline walking from muscle activity and walking mechanics. J Biomech. 2012;45(10):1842–9.

28. Russell DM, Apatoczky DT. Walking at the preferred stride frequency minimizes muscle activity. Gait Posture. 2016;45:181–6.

29. Jaramillo J. Are muscles tuned to the preferred stride frequency of walking? University of Massachusetts Amherst; 2020.

30. Prilutsky BI, Gregor RJ. Swing-and support-related muscle actions differentially trigger human walk–run and run–walk transitions. J Exp Biol. 2001;204(13):2277–87.

31. Stenum J, Choi JT. Neuromuscular effort predicts walk–run transition speed in normal and adapted human gaits. J Exp Biol. 2016;219(18):2809–13.

32. Segers V, Lenoir M, Aerts P, De Clercq D. Influence of M. tibialis anterior fatigue on the walk-to-run and run-to-walk transition in non-steady state locomotion. Gait Posture. 2007;25(4):639–47.

33. Neptune RR, Hull ML. A theoretical analysis of preferred pedaling rate selection in endurance cycling. J Biomech. 1999;32(4):409–15.

34. Marsh AP, Martin PE. The association between cycling experience and preferred and most economical cadences. Med Sci Sport Exerc. 1993;25(11):1269–74.

35. Vercruyssen F, Brisswalter J. Which factors determine the freely chosen cadence during submaximal cycling? J Sci Med Sport. 2010;13(2):225–31.

36. Rasmussen J, Damsgaard M, Voigt M. Muscle recruitment by the min/max criterion—a comparative numerical study. J Biomech. 2001;34(3):409–15.

37. Crowninshield RD, Brand RA. A physiologically based criterion of muscle force prediction in locomotion. J Biomech. 1981;14(11):793–801.

38. Redfield R, Hull ML. Prediction of pedal forces in bicycling using optimization methods. J Biomech. 1986;19(7):523–40.

39. Dingwell JB, Marin LC. Kinematic variability and local dynamic stability of upper body motions when walking at different speeds. J Biomech. 2006;39(3):444–52.

40. McDonald KA, Cusumano JP, Peeling P, Rubenson J. Multi-objective control in human walking: insight gained through simultaneous degradation of energetic and motor regulation systems. J R Soc Interface. 2019;16(158):20190227.

41. Stegeman D, Hermens H. Standards for surface electromyography: The European project Surface EMG for non-invasive assessment of muscles (SENIAM). 2007;

42. Peronnet F, Massicotte D. Table of nonprotein respiratory quotient: an update. Can J Sport Sci. 1991;16(1):23–9.

43. Kipp S, Byrnes WC, Kram R. Calculating metabolic energy expenditure across a wide range of exercise intensities: the equation matters. Appl Physiol Nutr Metab. 2018;43(6):639–42.

44. Devaprakash D, Weir GJ, Dunne JJ, Alderson JA, Donnelly CJ. The influence of digital filter type, amplitude normalisation method, and co-contraction algorithm on clinically relevant surface electromyography data during clinical movement assessments. J Electromyogr Kinesiol. 2016;31:126–35.

45. Kram R, Taylor CR. Energetics of running: a new perspective. Nature. 1990;346(6281):265–7.

46. Roberts TJ, Kram R, Weyand PG, Taylor CR. Energetics of bipedal running. I. Metabolic cost of generating force. J Exp Biol. 1998;201(19):2745–51.

47. Carrier D, Anders C, Schilling N. The musculoskeletal system of humans is not tuned to maximize the economy of locomotion. Proc Natl Acad Sci. 2011;108(46):18631–6.

48. Handsfield GG, Meyer CH, Hart JM, Abel MF, Blemker SS. Relationships of 35 lower limb muscles to height and body mass quantified using MRI. J Biomech. 2014;47(3):631–8.

49. Alexander RM. Optimization and gaits in the locomotion of vertebrates. Physiol Rev. 1989;69(4):1199–227.

50. Hunter LC, Hendrix EC, Dean JC. The cost of walking downhill: is the preferred gait energetically optimal? J Biomech. 2010;43(10):1910–5.

51. Rosenbaum DA, Gong L, Potts CA. Pre-crastination: hastening subgoal completion at the expense of extra physical effort. Psychol Sci. 2014;25(7):1487–96.

52. Kistemaker DA, Wong JD, Gribble PL. The central nervous system does not minimize energy cost in arm movements. J Neurophysiol. 2010;104(6):2985–94.

53. Westling G, Johansson RS. Factors influencing the force control during precision grip. Exp Brain Res. 1984;53(2):277–84.

54. Balasubramanian R, Howe RD, Matsuoka Y. Task performance is prioritized over energy reduction. IEEE Trans Biomed Eng. 2009;56(5):1310–7.

55. De Rugy A, Loeb GE, Carroll TJ. Muscle coordination is habitual rather than optimal. J Neurosci. 2012;32(21):7384–91.

56. Yandell MB, Zelik KE. Preferred Barefoot Step Frequency is Influenced by Factors Beyond Minimizing Metabolic Rate. Sci Rep. 2016;6:23243.

57. Monsch ED, Franz CO, Dean JC. The effects of gait strategy on metabolic rate and indicators of stability during downhill walking. J Biomech. 2012;45(11):1928–33.

58. Finley JM, Bastian AJ, Gottschall JS. Learning to be economical: the energy cost of walking tracks motor adaptation. J Physiol. 2013;591(4):1081–95.

59. Huang HJ, Ahmed AA. Older adults learn less, but still reduce metabolic cost, during motor adaptation. J Neurophysiol. 2014;111(1):135–44.

60. Whiting CS, Allen SP, Brill JW, Kram R. Steep (30°) uphill walking vs. running: COM movements, stride kinematics, and leg muscle excitations. Eur J Appl Physiol. 2020;120(10):2147–57.

61. Pontzer H. Economy and endurance in human evolution. Curr Biol. 2017;27(12):R613–21.

62. Bramble DM, Lieberman DE. Endurance running and the evolution of Homo. Nature. 2004;432(7015):345–52.

63. Noakes TDO. Fatigue is a brain-derived emotion that regulates the exercise behavior to ensure the protection of whole body homeostasis. Front Physiol. 2012;3:82.

64. Marino FE, Sibson BE, Lieberman DE. The evolution of human fatigue resistance. Journal of Comparative Physiology B, pp.1-12. J Comp Physiol B. 2022;192(3–4):411–22.

65. Proske U, Allen T. The neural basis of the senses of effort, force and heaviness. Exp Brain Res. 2019;237(3):589–99.

66. Monjo F, Terrier R, Forestier N. Muscle fatigue as an investigative tool in motor control: A review with new insights on internal models and posture–movement coordination. Hum Mov Sci. 2015;44:225–33.

67. Wong JD, O’Connor SM, Selinger JC, Donelan JM. Contribution of blood oxygen and carbon dioxide sensing to the energetic optimization of human walking. J Neurophysiol. 2017;118(2):1425–33.

68. Woledge RC, Curtin NA, Homsher E. Energetic aspects of muscle contraction. Monogr Physiol Soc. 1985;41:1–357.

69. Umberger BR, Gerritsen KG, Martin PE. A model of human muscle energy expenditure. Comput Methods Biomech Biomed Engin. 2003;6(2):99–111.

70. Beck ON, Punith LK, Nuckols RW, Sawicki GS. Exoskeletons improve locomotion economy by reducing active muscle volume. Exerc Sport Sci Rev. 2019;47(4):237–45.

71. Roberts TJ, Marsh RL, Weyand PG, Taylor CR. Muscular force in running turkeys: the economy of minimizing work. Science. 1997;275(5303):1113–5.

72. Berniker M, Kording K. Estimating the sources of motor errors for adaptation and generalization. Nat Neurosci. 2008;11(12):1454–61.

73. Taylor JL, Amann M, Duchateau J, Meeusen R, Rice CL. Neural contributions to muscle fatigue: from the brain to the muscle and back again. Med Sci Sport Exerc. 2016;48(11):2294.

